# Yeast two-hybrid screening identifies Dad1 as a candidate substrate of Hrr25 kinase in *Saccharomyces cerevisiae*

**DOI:** 10.1101/2025.07.29.667491

**Authors:** Meenakshi Agarwal, Santanu K. Ghosh, Sankalpa Chakraborty

## Abstract

In *Saccharomyces cerevisiae*, sister kinetochores are mono-oriented during meiosis I, ensuring that homologous chromosomes segregate to opposite poles, a process critically dependent on the kinase activity of Hrr25, a casein kinase. However, the direct substrates of Hrr25 involved in this mechanism remain poorly defined. In this study, we used a yeast two-hybrid (Y2H) approach to screen for and identify physical interactors of Hrr25. The *HRR25* gene was cloned into a Y2H bait vector, and its functional expression was confirmed by complementation of a temperature-sensitive *hrr25-ts* mutant. Screening independent Y2H libraries in three reading frames, followed by validation via reporter assays, restriction analysis, and sequencing, we identified six putative interactors: *HED1, DAD1, YDR015C* (from clone C1-5), *REP1* (C2-15), and *CYR1* and *SYS1* (C3-7). Phosphorylation site prediction and AlphaFold 3.0 structural modeling identified high-confidence Hrr25 target residues, including S70/T73 on Hed1 and S63 on Dad1, S323 on Rep1, and S198/S527 on Cyr1, each located in structurally accessible and potentially functional regions. Plasmid-dependent assays confirmed that reporter activation in C1-5 depended on the presence of the prey plasmid, and restriction mapping demonstrated that C1-5 contained a full-length *DAD1* ORF. Given Dad1’s known role in DASH/Dam1 kinetochore complex and its function in kinetochore-microtubule attachment, along with previous findings that *DAD1* mutations cause meiosis I defects, our data suggest that Dad1 may be a substrate of Hrr25. We propose that Hrr25-mediated phosphorylation of Dad1 could facilitate sister kinetochore co-orientation during meiosis I. These results provide new insights into the molecular mechanisms of chromosome segregation and identify Dad1 as a potential candidate substrate for Hrr25 in meiotic regulation.

## 1. Introduction

Accurate chromosome segregation during meiosis is essential for maintaining euploidy across generations. Errors in this process can have serious consequences, including aneuploidy, which may result in infertility, cancer, and congenital disorders such as Down syndrome ^1,2^. Similar to mitosis, meiotic chromosome segregation is controlled by the kinetochore, a large multi-protein complex that assembles at the centromere of chromosome and provides attachments sites for microtubules, thereby linking chromosome movements to microtubules dynamics^3–5^. However, meiosis involves additional layers of complexity and relies on four fundamental and conserved events: (1) homologous chromosomes pair and undergo recombination to form chiasmata, which physically link non-sister chromatids; (2) sister kinetochores of each homolog orient toward the same spindle pole (mono-orientation), enabling proper bi-orientation of homolog pairs on the meiosis I (MI) spindle; (3) cohesin is released in a stepwise manner during meiosis I and II to facilitate orderly chromosome segregation; and (4) Suppression of DNA replication prior to second meiotic division ^6^.

In *Saccharomyces cerevisiae* (*S. cerevisiae*), the mono-orientation of sister kinetochores during meiosis I is directed by the monopolin complex, comprising four proteins: Mam1, Csm1, Lrs4, and the kinase Hrr25. This complex is specifically recruited to the centromere during prophase I, where it promotes monopolar attachment of sister kinetochores to the spindle microtubules during meiosis I ^7,8^. Hrr25, a 494-amino acid multifunctional casein kinase I family member, has a well-characterized N-terminal kinase domain and a C-terminal proline/glutamine (P/Q)-rich region ^9^. It is essential for cell viability and exerts several functions during vegetative growth. Mutations in *HRR25* result in slow cell proliferation and increase sensitivity to DNA damage ^10^. During meiosis, a kinase-dead mutant of Hrr25 results in abnormal bi-orientation of sister chromatids at metaphase I and produces defective spores with unequal DNA content, highlighting the necessity of its kinase activity during meiosis I ^11^.

Despite its known significance, the full spectrum of Hrr25’s meiotic kinase substrates remain poorly understood. While Mam1, a component of the monopolin complex, has been identified as a phosphorylation target of Hrr25; however, this modification is dispensable for establishing mono-orientation during meiosis I ^11^. Additionally, Hrr25, in coordination with the Dbf4-dependent Cdc7 kinase, phosphorylates Rec8, a meiosis-specific cohesin subunit. This phosphorylation is required for Rec8 cleavage by separase during the transition from metaphase I to anaphase I ^12^. Nonetheless, there is currently no evidence that Rec8 phosphorylation by these kinases directly contributes to the establishment of sister chromatid mono-orientation.

These findings suggest that other kinetochore or chromatin-associated proteins may serve as critical Hrr25 substrates, whose phosphorylation may be essential for promoting sister chromatid mono-orientation in meiosis I. Identifying these targets is therefore crucial for uncovering the molecular mechanisms governing accurate meiotic chromosome segregation. We further hypothesize that Hrr25 must physically associate with its substrates at the kinetochore to exert its kinase activity. To test this, we employed a yeast two-hybrid system using Hrr25 as bait to screen a yeast genomic library. This approach enabled the identification of candidate Hrr25-interacting proteins, which may represent functionally relevant kinase substrates and offer new mechanistic insights into the regulation of meiotic chromosome segregation.

## 2. Material and methods

### Yeast strains, bacterial strains, plasmids, and primers

All the yeast strains used and constructed in this study are specified in Table S1. All the bacterial strains used in this study are specified in Table S2. All the plasmids used and constructed in this study are listed in Table S3. The primers used in this study are listed in Table S4.

### Growth media and growth condition

Yeast strains were cultured on YPD medium (1% yeast extract, 2% peptone, 2% dextrose) at 30°C unless otherwise specified. For screening of transformants, synthetic dropout (SD) media containing yeast nitrogen base (YNB), dextrose, and dropout supplements lacking specific amino acids, as required by the experiment, were used. Bacterial strains, either transformed with plasmids or untransformed, were grown in Luria-Bertani (LB) broth or on LB agar plates at 37°C.

### Construction of recombinant plasmids

Plasmid DNA isolation, preparation of competent *E. coli* cells, and bacterial transformation procedures were carried out manually following protocols described in *Molecular Cloning: A Laboratory Manual* by Green and Sambrook, 4th edition (Cold Spring Harbor Laboratory Press).

To construct the recombinant bait plasmid, total genomic DNA was isolated from yeast cells and used as a template to PCR amplify the *HRR25* gene. Specific restriction enzyme restriction sites, *BamH*1 and *Pst*1, were incorporated into the forward and reverse primers. The plasmid vector DNA, pGBDC1 was isolated from *E. coli*, digested with the same restriction enzymes used for the PCR product, and subsequently ligated to the *HRR25* insert. The ligation mixture was transformed into competent *E. coli* cells. Transformants were screened using a combination of assays including agarose gel electrophoresis to assess plasmid size shift (retardation assay), restriction enzyme digestion, and PCR amplification of the insert.

### Amplification and quality assessment of yeast two-hybrid library DNA

Yeast two-hybrid (Y2H) libraries carried in the *pGAD* vector, representing three different reading frames (Y2HL-C1, Y2HL-C2, Y2HL-C3), were used for transformation. Competent *E. coli* DH5α cells were individually transformed with each *pGAD* vector carrying a library in a specific reading frame. The transformed cells were cultured, and plasmid DNA was extracted using a maxi-prep protocol to amplify the libraries.

To assess the quality and efficiency of the amplified libraries, a small aliquot of each plasmid preparation was transformed into competent *E. coli* cells. Plasmid DNA was then isolated from at least 10 individual colonies per library and subjected to restriction enzyme digestion using *SmaI* and *PstI*. Library efficiency was calculated as the percentage of plasmids containing an insert relative to the total number analyzed. The observed efficiency was approximately 80% for Y2HL-C1 and Y2HL-C2, and 85% for Y2HL-C3. To ensure comprehensive coverage, the amplified yeast two-hybrid (Y2H) library should represent the yeast genome approximately three times. The number of yeast colonies required for screening was calculated using the following formula:

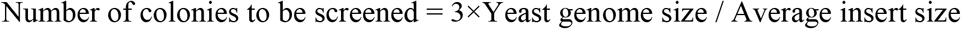

### Yeast two-hybrid studies

Y2H analyses were carried out using the *S. cerevisiae* strain PJ69-4A, which contains three reporter genes, *HIS3, ADE2*, and *LacZ*, used to detect protein-protein interactions. Plasmids pGBD and pGAD, with or without gene fusions, were transformed into PJ69-4A. For library screening, the PJ69-4A strain harboring the bait construct (pGBD vector with *TRP1* marker and gene insert, *HRR25*) was transformed with a *S. cerevisiae* genomic two-hybrid library cloned into a pGAD vector carrying the *LEU2* selection marker ^13^. Transformants were selected on SD dropout plates lacking leucine and tryptophan (SD/-Leu-Trp) to maintain both plasmids.

Positive interactors were first screened by patching colonies onto SD/-Leu-Trp-His plates supplemented with 1 mM 3-amino-1,2,4-triazole (3-AT) to suppress background *HIS3* expression. Colonies that grew on this medium were further tested on SD/-Leu-Trp-Ade plates. Those showing growth on both selective mediums were subjected to β-galactosidase (β-Gal) assays to confirm *LacZ* reporter activation.

To recover library plasmids from positive clones, total yeast DNA was isolated and transformed into *E. coli* KC8 cells. Transformants were selected on minimal medium lacking leucine, allowing recovery of the *pGAD*-based library plasmids.

### Colony-lift filter assay for β-galactosidase activity

To detect β-galactosidase activity, yeast colonies were patched onto SD selection plates alongside appropriate positive and negative control strains. After full colony growth, a circular nitrocellulose (NC) filter matching the size of the Petri dish was gently placed over the colonies and allowed to sit for 2–3 minutes to transfer cells. The NC filter with the adhered colonies was then carefully removed and placed in a separate Petri dish, followed by rapid freezing using liquid nitrogen poured directly onto the filter. Once frozen, the filter was transferred to a fresh Petri dish pre-lined with Whatman filter paper. An aliquot of Z buffer (containing Na_2_HPO_4_·7H_2_O, NaH_2_PO_4_, KCl, MgSO_4_·7H_2_O, β-mercaptoethanol, and distilled water) supplemented with X-gal at a final concentration of 1 mg/mL was gently added from the edge to saturate the NC filter. The setup was incubated at 30°C for 1–2 hours. Colonies exhibiting a visible blue or purple color change were considered positive for β-galactosidase activity.

### Recovery of plasmid DNA from yeast cells

To recover library plasmids from yeast, total genomic DNA was isolated from a 5 mL overnight grown yeast culture. The DNA was resuspended in 50 µL of TE buffer. An aliquot of 10 µL was then transformed into competent *E. coli* KC8 cells. Transformants were selected on LB agar plates supplemented with ampicillin. Plasmid DNA was subsequently isolated from bacterial colonies and analyzed for the presence of inserts via restriction enzyme digestion.

### Plasmid dependent yeast two-hybrid assay

To confirm that reporter activation was dependent on both bait and prey plasmids, a plasmid-dependent assay was performed. A library-derived yeast transformant that tested positive for all three reporter genes (*HIS3, ADE2*, and *LacZ*) was selected. This strain, containing both the bait (*TRP1* marker) and prey (*LEU2* marker) plasmids, was initially grown on SD/-Leu/-Trp medium and then cultured in YPD broth for approximately 40 generations to allow plasmid segregation. Following growth, cells were streaked onto YPD plates to isolate single colonies. Approximately 40–50 single colonies were patched onto YPD, SD/-Leu, and SD/-Trp plates. All colonies were expected to grow on YPD. Colonies that showed growth on SD/-Trp but failed to grow on SD/- Leu were identified as having retained only the bait plasmid (Trp^+^, Leu^−^) and lost the prey plasmid. These selected colonies, now lacking the prey plasmid, were expected to lose reporter activity, thereby confirming that interaction-driven reporter activation required the presence of both plasmids. Reintroduction of the prey plasmid into these strains should restore reporter activity, further validating the interaction.

### Bioinformatic Analyses of potential Hrr25 Phosphorylation sites on target Proteins

*S. cerevisiae* specific bioinformatic tools (NetPhosYeast 1.0 ^14^, GPS 6.0 ^15^) and non-species-specific (NetPhos-3.1 ^16^, PhosphoSVM ^17^) were used for sequence-based phosphorylation site predictions. For both species specific and non-specific tools used in this study, the target sites were selected and sorted for Casein Kinase 1 (CK1) and the parameters were kept as default. The amino acid sequences for the Hrr25 and potential target proteins were retrieved from *Saccharomyces* genome database (SGD): Hrr25 (SGD: S000006125), Hed1 (SGD: S000113613), Cyr1 (SGD: S000003542), YDR015C (SGD: S000002422), Rep1 (SGD: S000029675), Sys1 (SGD: S000003541), Dad1 (SGD: S000002423). The FASTA sequences of the target proteins were used as the inputs for phosphorylation site analyses. The outputs from all the tools were manually compared to identify the common phosphorylation sites on Hrr25 target proteins.

Due to the unavailability of complexed structures in protein databank (PDB), AlphaFold 3.0 ^18^ was used to generate the complex structures of Hrr25 target proteins. The FASTA sequences from SGD were used as inputs in AlphaFold 3.0 with the default parameters. The top ranked complexed structures for each interaction were used for final processing in PyMol for better visualization and image generation.

## 3. Results

### 3.1 Recombinant HRR25 bait plasmid complemented temperature sensitivity in hrr25-ts yeast

The *HRR25* open reading frame (ORF) was PCR-amplified from wild-type *S. cerevisiae* genomic DNA using primers MA53 and MA54 (Table S4), and cloned into the pGBD-C1 vector using *BamHI* and *PstI* restriction sites. The resulting recombinant plasmid, designated pGBDC1-*HRR25* (pSGB96), was validated by restriction enzyme digestion, which produced the expected ∼1.5 kb *HRR25* insert fragment along with the vector backbone (Figure 1A-C).

**Figure 1.**
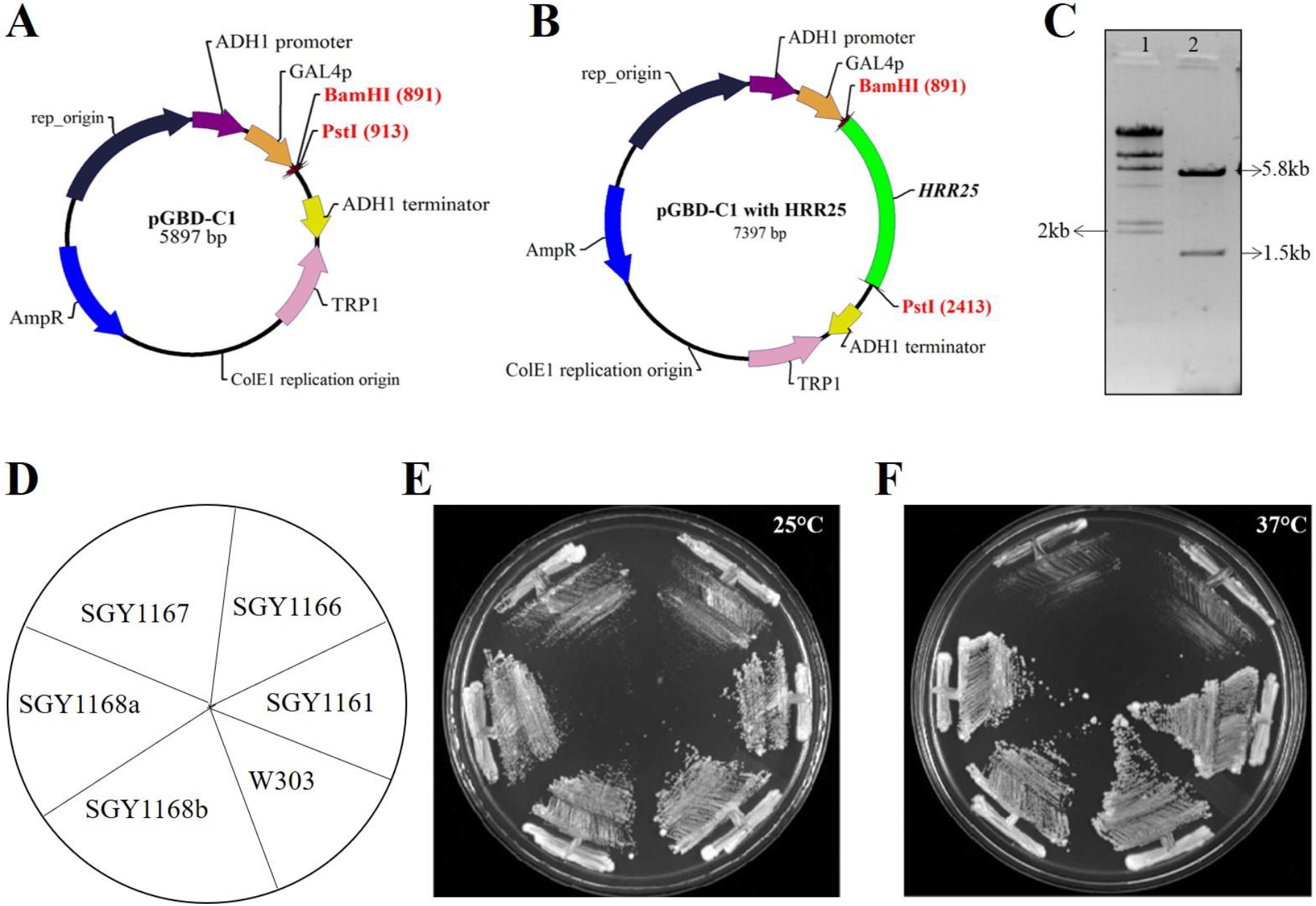
Construction and functional validation of bait vector. *Cloning of HRR25 in pGBD-C1* (**A-C**). The vector map of pGBD-C1 (**A**), Vector map of pGBDC1 with cloned *HRR25* ORF (**B**). Cloned fragment is shown in green color. RE digestion of clone with *Bam*HI and *Pst*I (**C**). Lane 1, λ *Hin*dIII marker DNA, Lane 2, digested product. *pGBD-C1 carrying HRR25 ORF complemented the growth in hrr25-ts mutant* **(D-F)**. Position of strains on plate (**D**). SGY1168 a and b represent two different clones, Cells were streaked on YPD plates with appropriate controls and incubated at 25°C (**E**) and 37°C (**F**). The photograph was taken after two days of growth.

To confirm functional expression of the bait construct, a complementation assay was performed using a temperature-sensitive *hrr25-ts* mutant strain. This strain (SGY1166) grows optimally at 25 °C but exhibits impaired or no growth at 37 °C. Transformation of the *hrr25-ts* mutant with pGBDC1-*HRR25* (SGY1168a, b) successfully restored growth at the non-permissive temperature of 37 °C, indicating that the cloned *HRR25* gene is functionally expressed. In contrast, cells transformed with the empty vector (SGY1167) or left untransformed (SGY1166) failed to grow under these conditions, confirming the specificity of the complementation (Fig. 1D-F).

### 3.2 Yeast Two-Hybrid Library Screening resulted in six putative interactors of Hrr25

The host yeast strain PJ69-4A carrying bait plasmid pSGB96 and three dependent yeast two-hybrid (Y2H) libraries were screened individually. A known protein–protein interaction between kinetochore proteins Chl4 and Mcm19 ^19^ was used as a positive control. The AD-Chl4 and BD-Mcm19 combination activated all three reporters, whereas negative controls (empty vectors) showed no reporter activity (Figure 2). Across all three libraries, 19 transformants tested positive for all three reporters, SD/-Trp/-Leu/-His with 1 mM 3-AT, SD/-Trp/-Leu/-Ade, and β-Gal assays. These positive clones were subjected to plasmid recovery as described in Section 2. Prey plasmids were isolated, and inserts were analyzed by restriction enzyme digestion. Clones containing inserts larger than 500 bp were selected for further analysis, yielding six candidates. Of these, six clones exhibiting strong reporter activation (Figure 2) sequenced using a primer specific to the Gal4 activation domain region of the vector. Sequence data were obtained only from three clones and used to perform BLAST searches against the *S. cerevisiae* genome database. The resulting genomic coordinates and corresponding ORFs are listed in Table 1.

**Figure 2.**
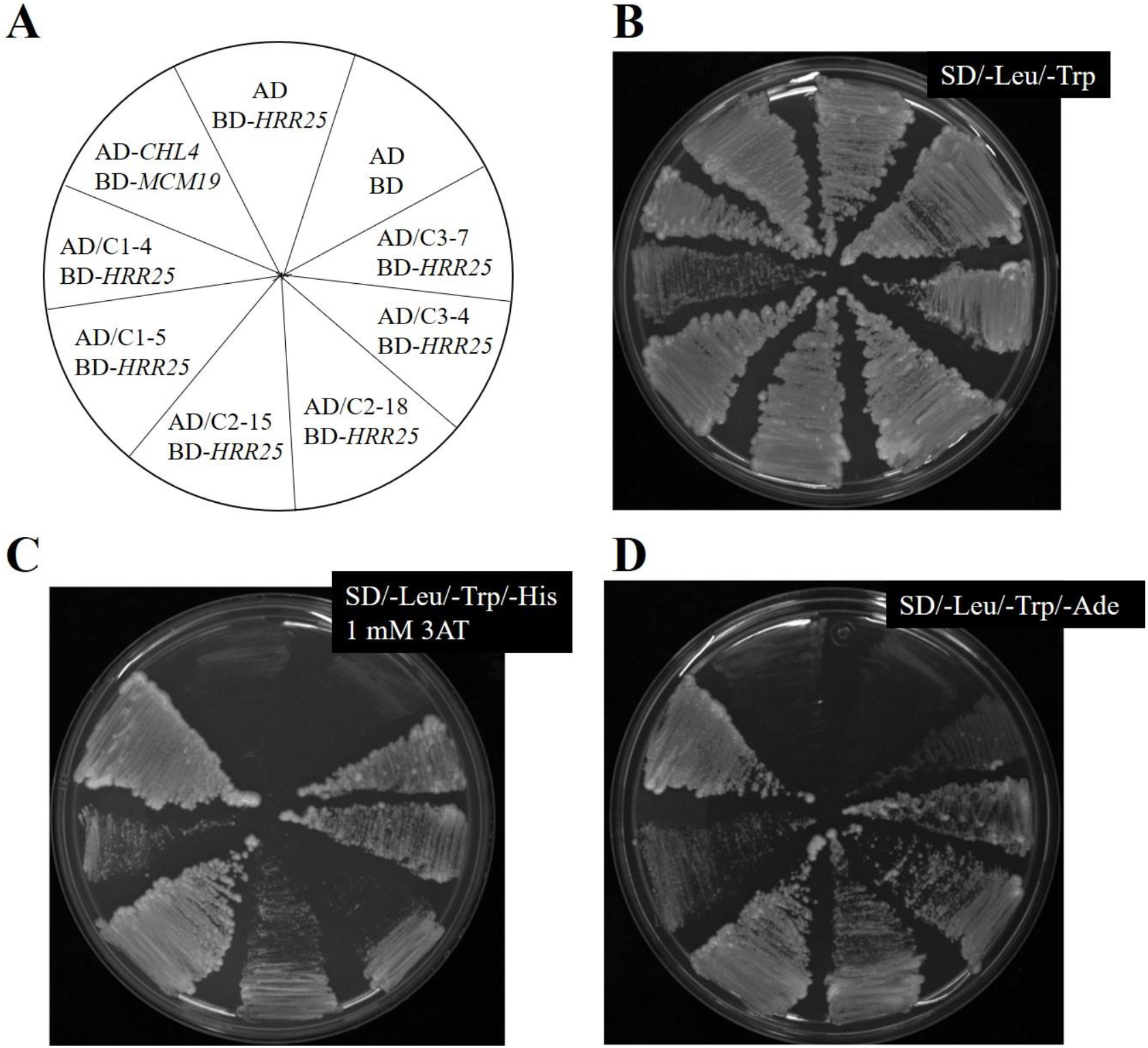
Reporter assay for the putative interactors. A schematic diagram showing name and position of strains on the plate (**A**). AD-*CHL4* BD-*MCM19* was taken as a positive control. AD BD-*HRR25* and AD BD were used as a negative control. C1, C2 and C3 denote transformants of Y2HL-C1, Y2HL-C2 and Y2HL-C3 libraries. Cells were streaked on SD/-Leu/-Trp plate **(B)**, SD/-Leu/-Trp/-His dropout plate with 1mM 3AT **(C)**, and SD/-Leu/-Trp/-Ade plate **(D)**.

**Table 1.** Summary of BLAST result from query sequences against *S. cerevisiae* genome.

| **Name of clone** | **Identified coordinates from BLAST result** | **Identified ORFs within the coordinates** | **Homology (bps)** |
| --- | --- | --- | --- |
| C1-5 | 478021...478579 | *HED1*(Chr. IV 477797...478285) | 266 |
|  |  | *DAD1*(Chr IV 478474...478758) | 108 |
|  |  | *YDR015C* (Chr. IV 477960...478199) | 180 |
| C2-15 | 1606...2531 | *REP1* (1887...3008) | 648 |
| C3-7 | 430767...431361 | *CYR1* (Chr X 425157...431237) | 474 |
|  |  | *SYS1* (Chr X- 431589...432200) | 15 |

### 3.3 Structural modeling identified the phosphorylation sites on putative target proteins

Further phosphorylation sites were predicted on the putative Hrr25 target proteins. To identify conserved serine (S) and threonine (T) residues likely to be phosphorylated by Hrr25, we employed multiple bioinformatic tools for phosphorylation site prediction. S/T residues consistently predicted by at least two tools were considered potential Hrr25 phosphorylation sites. To prioritize high-confidence candidates, we examined the spatial proximity and orientation of these predicted residues within Hrr25 target protein complex models generated using AlphaFold 3.0.

Based on structural analysis, including distance and positioning, several predicted sites were identified as high-confidence phosphorylation targets (listed in Table 2). We identified two high-confidence phosphorylation sites each on Hed1 and Cyr1, and one site each on Dad1 and Rep1. In Hed1, both S70 and T73 were consistently predicted and found in close proximity, suggesting their accessibility for phosphorylation by Hrr25. On Dad1, S63 was appropriately positioned and oriented for potential phosphorylation, whereas T34 appeared distant and poorly oriented, making phosphorylation by Hrr25 less likely. For Cyr1, two high-confidence sites S198 and S527 were identified. Although additional sites such as S325 and S582 were predicted with strong scores by three tools, their spatial positioning was unfavorable for Hrr25-mediated phosphorylation. Rep1 contained one high-confidence site, S323. Several of these high-confidence sites, including T73 on Hed1, S198 and S527 on Cyr1, and S323 on Rep1, were located in surface-exposed loops or flexible regions, which could facilitate access by Hrr25. In contrast, the predicted sites on Sys1 and YDR015C were found to be spatially distant or poorly oriented, making them unlikely targets for Hrr25 phosphorylation (Figure 3).

**Table 2.**
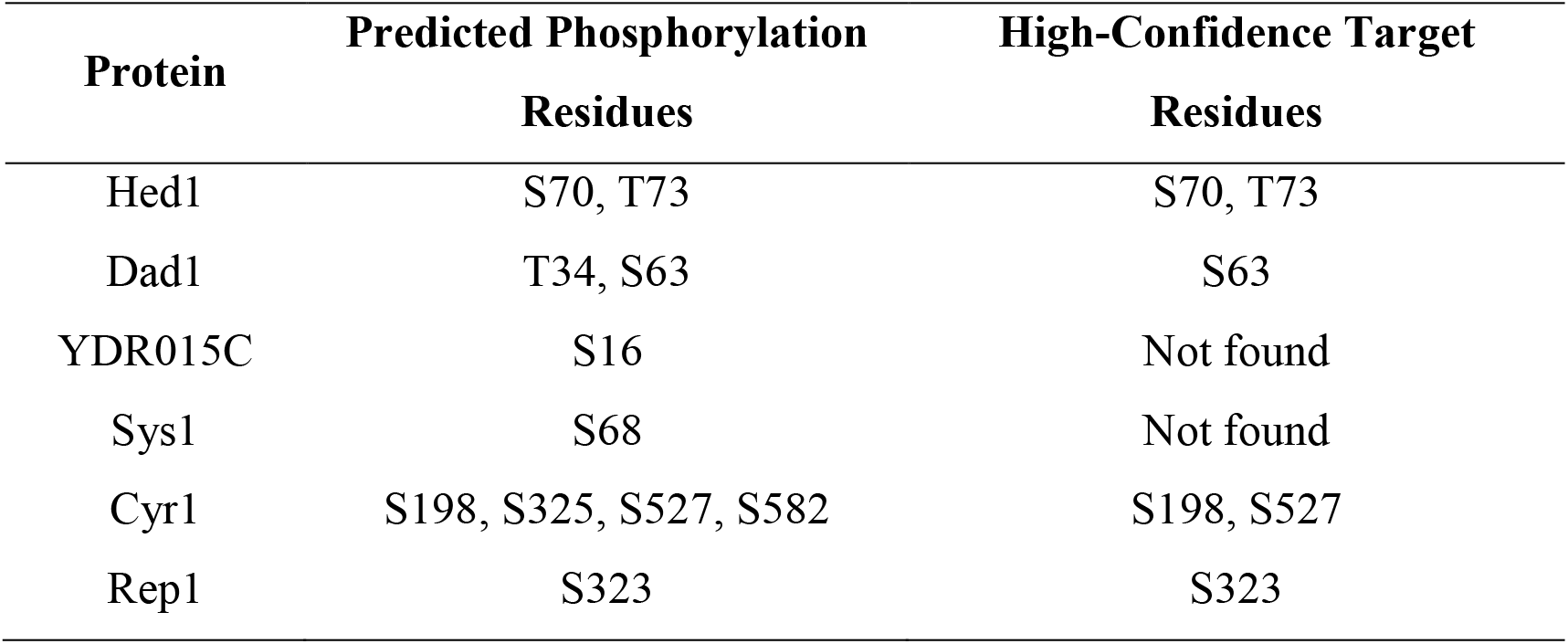
Table summarizes the predicted phosphorylation and high-confidence target residues for each protein listed.

**Figure 3.**
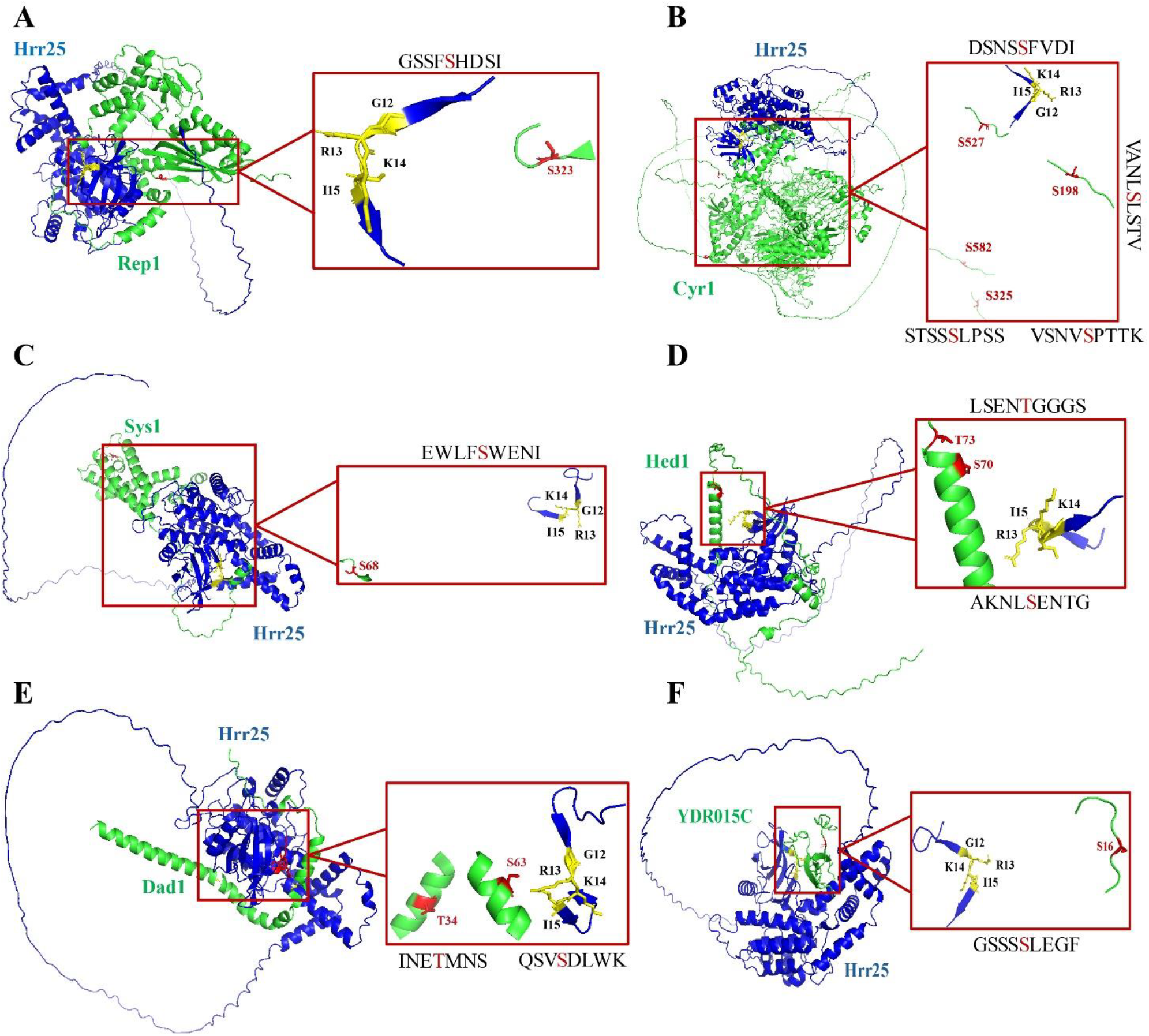
Structural model showing predicted phosphorylation sites on potential Hrr25 interacting target proteins. Panels A-F display the docked complexes of Hrr25 (blue) with its putative target proteins (green): A) Rep1, B) Cyr1, C) Sys1, D) Hed1, E) Dad1 and F) YDR015C. Predicted phosphorylation sites on the target proteins are shown in red. Insets zoom in on the interaction interface, illustrating the spatial arrangement of key catalytic residues in Hrr25 (yellow) relative to the predicted phosphorylatable serine/threonine residues (red) on the targets.

### 3.4 Reporter expression was found linked to the presence of library plasmid in C1-5

To further validate the positive interaction between Hrr25 and the putative target proteins, a plasmid-dependent assay was performed as described in Section 2. Our sequence analysis of the encoded proteins revealed that clone C2-15 carried the Rep1 protein, which originates from the 2-micron plasmid (Table 1). Because Rep1 is non-chromosomal and unlikely to be involved in chromosome segregation, C2-15 was excluded from further investigation. Only transformants C1-5 and C3-7 were used in subsequent experiments. The colony obtained from C1-5 transformant, bearing the pGBD-*HRR25* plasmid but lacking the prey plasmid (Trp^+^ Leu^-^), did not show any reporter assay and behaved similarly to the negative control (Figure 4). However, when this strain was re-transformed with the prey plasmid (Trp^+^ Leu^+^), reporter expression including β-Gal assay was restored, indicating a rescued interaction (Figure 4). In contrast, the C3-7 transformant tested negative for the adenine reporter in the plasmid-dependent assay (Figure 4D). However, Positive control strains carrying pAD-Chl4/pBD-Mcm19 were streaked on all plates and consistently showed reporter activity, validating the assay conditions.

**Figure 4.**
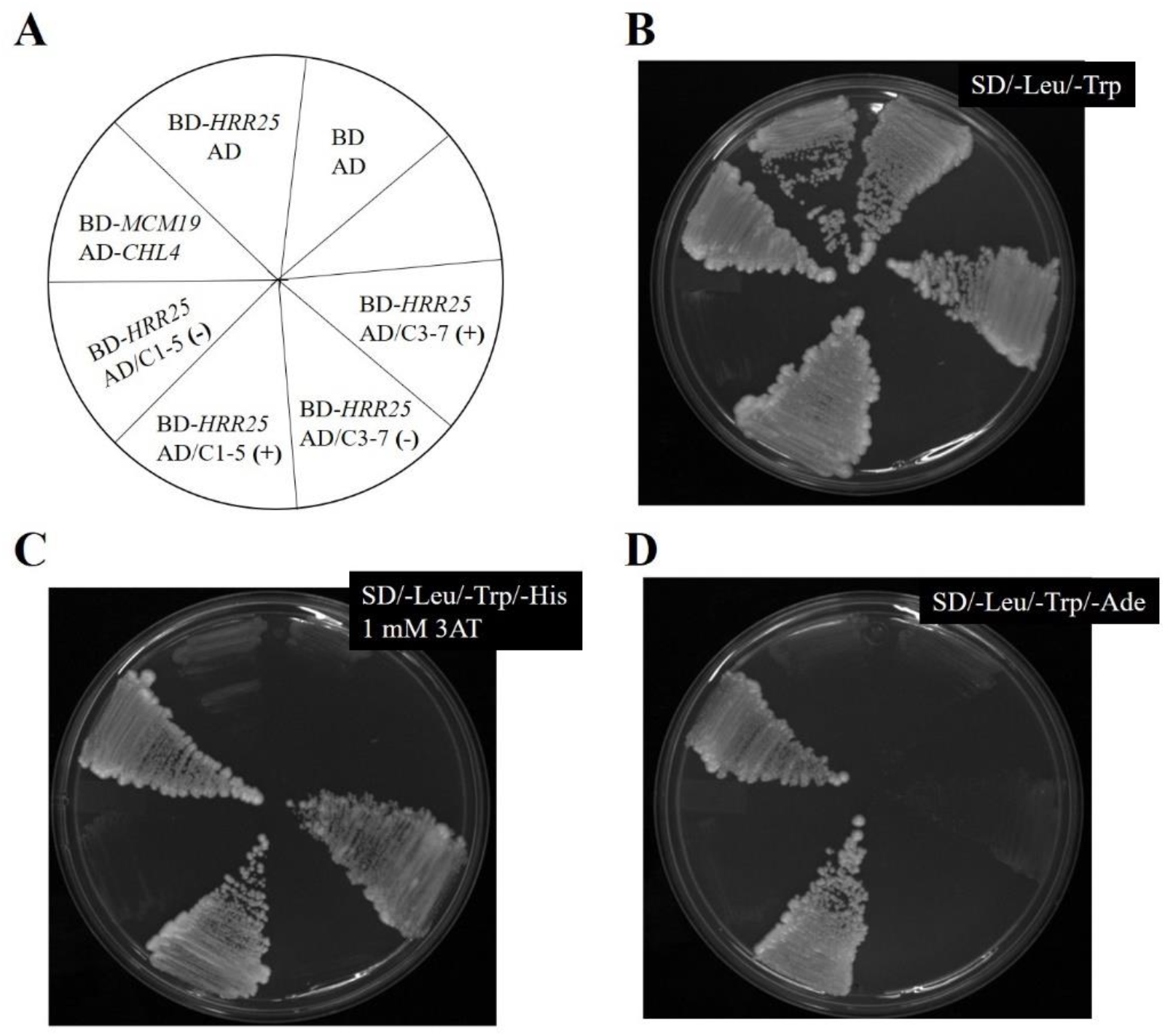
Plasmid dependent assay from transformants, C1-5 and C3-7. A schematic diagram showing name and position of strains on the plate (**A**). AD-*CHL4* BD-*MCM19* was taken as a positive control. AD BD-*HRR25* and AD BD were used as a negative control. Transformants C1-5 and C3-7 were tested under both prey plasmid-loss (-) and retransformation (+) conditions. Cells were streaked on Leu and Trp dropout plate **(B)**, SD/-Leu/-Trp/-His with 1mM 3 AT plate **(C)**, and SD/-Leu/-Trp/-Ade plate **(D)**.

### 3.5 Restriction mapping revealed a full-length DAD1 ORF within the C1-5

Since sequencing was conducted using primer targeting only the Gal4 activation domain, it did not cover the entire insert. To determine the complete extent of the insert present in the C1-5 transformant, restriction mapping was carried out (Figure 5A). This analysis revealed that the plasmid contained truncated versions of the *HED1* and *YDR015C* ORFs, as well as a full-length *DAD1* ORF (Figure 5B).

**Figure 5.**
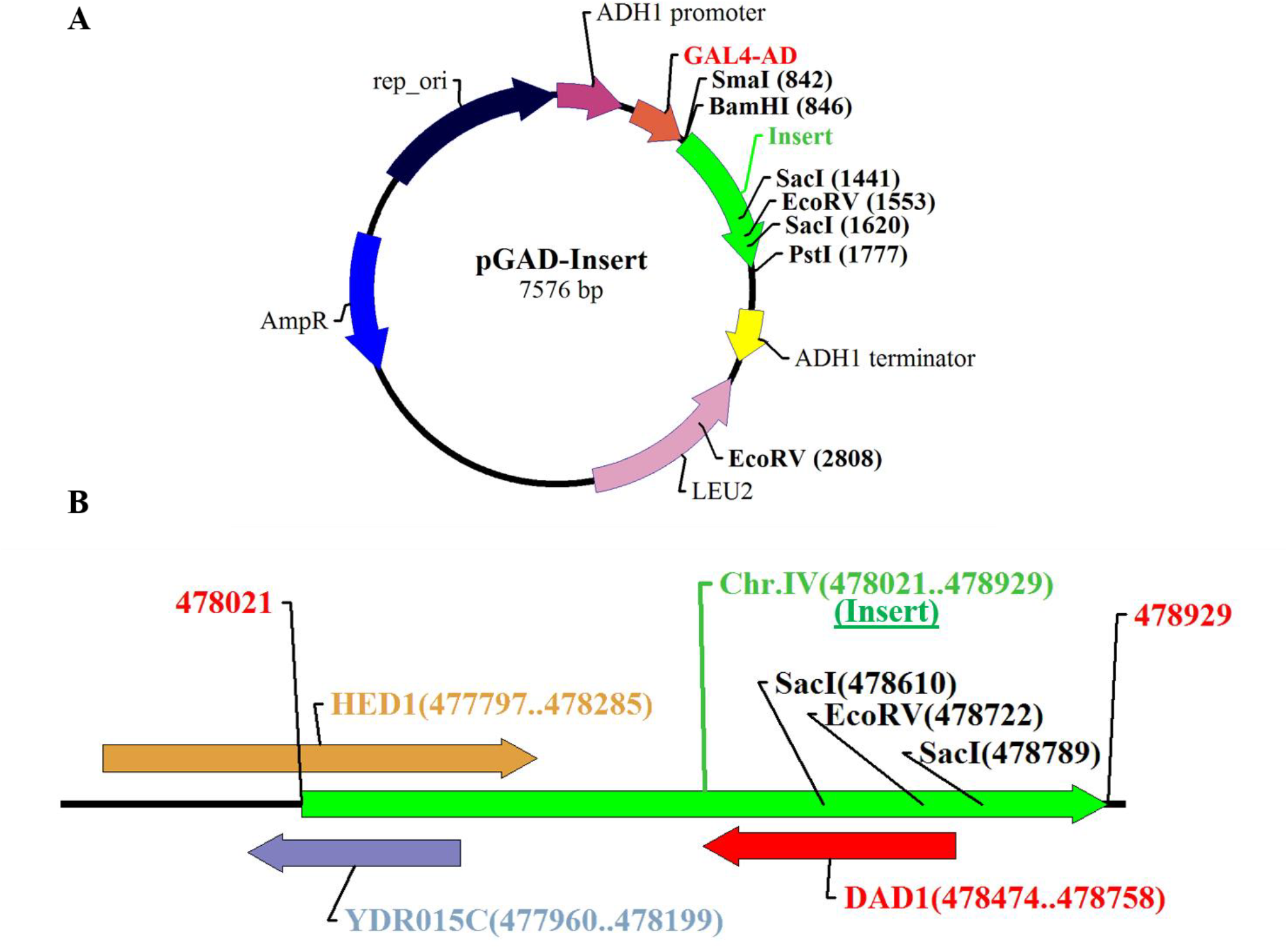
Restriction mapping of the insert present in C1-5. **(A)** Vector map of pGAD-C1 plasmid carrying insert. The insert is shown in green color. The position of restriction enzymes is marked on the insert. **(B)** Linearized restriction map of the insert **(B)**. Position of ORFs and their coordinates are shown. Each ORF is represented in different color.

## 4. Discussion

Given that the kinase activity of Hrr25 is critical for monopolin function at the centromere, this study aimed to identify potential substrates of Hrr25 that could help elucidate the mechanism underlying chromosome segregation during meiosis I. To achieve this, we employed a yeast two-hybrid system in conjunction with genetic approaches to screen for Hrr25-interacting proteins.

We successfully amplified yeast two-hybrid genomic libraries in all three reading frames with high transformation efficiencies. The functional integrity of the BD-Hrr25 fusion protein was confirmed via complementation assays, and the screening system was validated using established positive and negative controls. This setup allowed us to recover and analyze 19 initial positive clones, which were narrowed down to six clones through restriction digestion and further to three, C1-5, C2-15, and C3-7 via sequencing.

Among the three sequenced clones, clone C2-15 contained an insert encoding Rep1, a protein involved in the partitioning of the 2μm plasmid during mitosis ^20^. Clone C3-7 included two open reading frames, *CYR1* and *SYS1*, which encode adenylate cyclase and a Golgi-resident membrane protein, respectively ^21,22^. Given their known cellular roles and lack of kinetochore localization, these proteins are unlikely to be involved in chromosome segregation. Although phosphorylation prediction models indicated potential casein kinase target sites in these proteins, the lack of biological relevance to kinetochore function led to their exclusion. Clone C1-5 was further evaluated through plasmid-dependent assays to confirm that the observed reporter activation was specifically due to the presence of the plasmid insert. These assays supported the initial positive result, validating clone C1-5 for subsequent characterization.

### Is Hrr25-mediated phosphorylation of Dad1 important for meiosis I?

Restriction mapping analysis on insert obtained from clone C1-5 showed the presence of three open reading frames (ORFs): *HED1, DAD1*, and *YDR015C*. Of these, complete ORF of *DAD1* was present whereas ORFs belonging to *HED1* and *YDR015C* were presented in truncated form although in-frame fusion with the GAD sequences was maintained.

*HED1* encodes Hed1, a meiosis-specific protein known to regulate meiotic recombination by inhibiting the Rad51 recombinase. Previous studies have established that Hed1 is phosphorylated by the meiosis-specific kinase Mek1, particularly at T40, to modulate Rad51 activity. Additional phosphorylation sites at S38 and S42, located at N-terminus, have also been identified via mass spectrometry ^23^. Moreover, our in-silico model predicted potential phosphorylation sites at S70 and T73 and also the accessibility for phosphorylation by Hrr25. However, all of these residues lie outside the truncated region that captured in our screen. To our knowledge, there is no literature indicating a predicted interaction between Hed1 and Hrr25, nor is Hed1 known to be a substrate of Hrr25. Therefore, we consider it is less likely that Hed1 is a target of Hrr25. Nonetheless, to conclusively rule out this possibility, further analyses such as full-length interaction assays and co-immunoprecipitation are needed.

Another detected full length ORF belongs to Dad1 protein. Dad1 protein localizes to the kinetochore and an essential subunit of the DASH/Dam1 complex in budding yeast. This complex is involved in kinetochore-microtubule attachment during chromosome segregation in both mitosis and meiosis ^24^. The Dam1 complex is known to be regulated by phosphorylation; in mitosis, its subunits are known to be phosphorylated by the Ipl1 and Mps1, and Cdk1 kinases ^25^. However, direct phosphorylation of Dad1 by Ipl1 kinase has not been demonstrated ^25–28^.

In meiosis, Dam1 complex components are also subjected to phosphoregulation by kinase Mps1 which promotes chromosome biorientation. However, Mps1 inactivation does not produce major catastrophic effect during meiosis when Dam1 is not phosphorylated by Mps1 ^29^. Importantly, a recent study reported that *DAD1*^E50D^ mutation disrupted meiosis I-specific chromosome segregation and this mutation lies within a conserved phosphorylation motif predicted to be targeted by Hrr25 ^30^. Our phosphorylation prediction model also identified S63 on Dad1, with both its position and orientation favoring potential phosphorylation. Given that both Hrr25 and Dad1 localize to kinetochore in meiosis, and in light of both our result and existing literature, we propose that Dad1 may be a substrate of Hrr25. This interaction may facilitate sister kinetochore co-orientation during meiosis I as illustrated in Figure 6. Further, confirmatory experiments such as co-immunoprecipitation, targeted yeast two-hybrid assays, and phospho-mutant functional analyses are needed for next steps. It would be interesting to investigate whether phosphorylation of Dad1 by Hrr25 is required for the transition from meiosis I to meiosis II and for faithful chromosome segregation.

**Figure 6.**
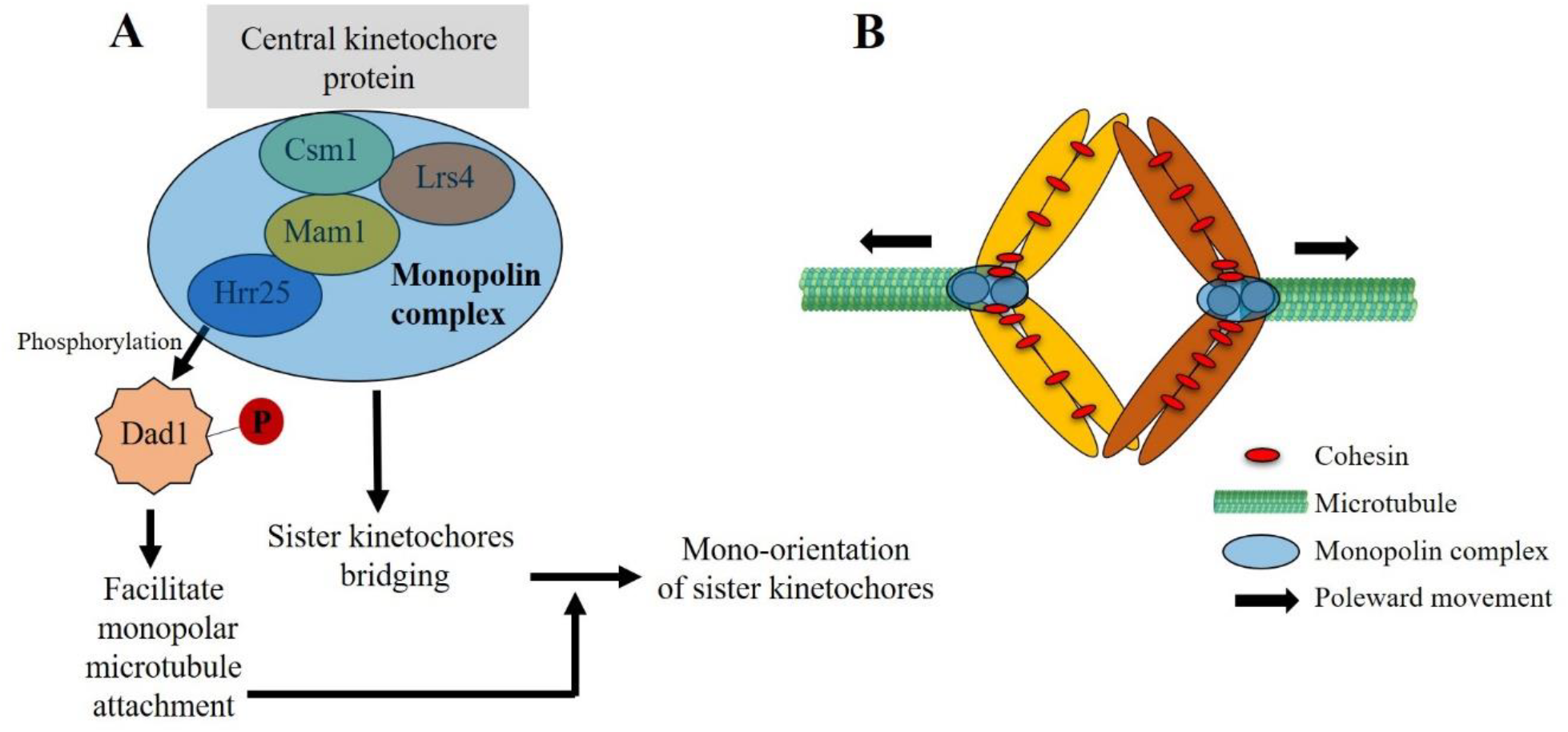
Model showing the role of Dad1 phosphorylation by Hrr25 during meiosis I in *S. cerevisiae*. **(A)** The monopolin complex facilitates sister kinetochores clamping, while Hrr25 kinase phosphorylates Dad1, promoting oligomerization or stabilization of the Dam1 complex around the microtubule. These coordinated events ensure the mono-orientation of sister kinetochores. **(B)** As a result, homologous chromosomes achieve biorientation and undergo directed movement from metaphase I to anaphase I.

In addition to *HED1* and *DAD1*, the insert in clone C1-5 also contained the dubious ORF *YDR015C*, whose function remains uncharacterized. Our phosphorylation prediction model identified a potential casein kinase target site at S16 within the region captured in our screen, although the confidence for this prediction was low. While speculative, *YDR015C* may encode a previously unrecognized Hrr25 substrate. Further validation through pairwise yeast two-hybrid interactions would be required to confirm its interaction.

It is also important to note that we do not rule out the possibility of additional physical interactors of Hrr25 that may have gone undetected due to limitations of the yeast two-hybrid system or other aspects of our experimental design. In addition, our study used a combination of bioinformatic phosphorylation prediction tools and AlphaFold 3.0 to identify the potential Hrr25 phosphorylation sites on the target proteins. While the accuracy of AlphaFold 3.0 in predicting the complexed protein structures is high, the potential changes and variability in the molecular dynamics during enzyme-substrate reaction might alter the structures of the interacting proteins. This in turn can change the orientations of the target proteins, rendering the positioning of the phosphorylation sites. Our approach can overlook this possibility, which can be considered a limitation of this section of the study. Future work incorporating molecular dynamics simulations or experimental validation would help address this limitation and provide a more comprehensive understanding of phosphorylation site accessibility in vivo.

## Conclusion

Overall, this study provides preliminary but compelling evidence for potential Hrr25 substrates. The identification of Dad1 as a candidate substrate is particularly intriguing given its critical role in chromosome segregation and the recent evidence linking it to meiotic defects. Likewise, the interaction between Hed1 and Hrr25, if validated, could reveal a novel layer of meiotic recombination regulation. Further biochemical and genetic investigations will be essential to substantiate these findings and fully elucidate the interactor of Hrr25 in meiosis. The result lay the groundwork for future investigations that could uncover novel mechanisms governing chromosome segregation during meiosis.

## Authors contribution

MA contributed to the conceptualization, design, data curation, funding acquisition, investigation, methodology, resources, software, supervision, validation, visualization, writing-original draft, writing-review and editing. SKG contributed to conceptualization, investigation, funding acquisition, resources, and supervision. SC contributed to data curation, methodology and software. All authors read and approved the final manuscript.

