## Supplemental File for "Yeast two-hybrid screening identifies Dad1 as a candidate substrate of Hrr25 kinase in *Saccharomyces cerevisiae*"

**Supplementary information:**

**Table S1**. List of yeast strains constructed in this study.

Parent strain of W303 with the following genotype was used to construct the strains:

*MAT***a** *ura3-52, lys2-801, ade2-101, trp1-*Δ*63, his3-*Δ*200, leu2-*Δ*1*

| **Strain** | **Mating type** | **Genotype** |
| --- | --- | --- |
| SGY1070 | *MAT***a** | *trp1-901 leu2-3, 112 ura3-52 his3-200 gal4*Δ *gal80*Δ *Met2::GAL7-lacZ LYS2::GAL1-HIS3 GAL2-ADE2 pGBDC1:: TRP1 + pGAD424::LEU2* |
| SGY1158 | MAT**a** | *trp1-901 leu2-3, 112 ura3-52 his3-200 gal4*Δ *gal80*Δ *Met2::GAL7-lacZ LYS2::GAL1-HIS3 GAL2-ADE2 pGBDC1-HRR25::TRP1* |
| SGY1161 | MAT**a** | *trp1-901 leu2-3, 112 ura3-52 his3-200 gal4*Δ *gal80*Δ *Met2::GAL7-lacZ LYS2::GAL1-HIS3 GAL2-ADE2 pGBDC1:: TRP1* |
| SGY1166 | *MAT***a** | *ura3Δ0 leu2Δ0 his3Δ1 lys2Δ0 met15Δ0 can1Δ0::LEU2-MFA1pr::HIS3, hrr25-ts::URA3* |
| SGY1167 | *MAT***a** | *ura3Δ0 leu2Δ0 his3Δ1 lys2Δ0 met15Δ0 can1Δ0::LEU2-MFA1pr::HIS3, hrr25-ts::URA3 + pGBDC1:: TRP1* |
| SGY1168a | *MAT***a** | *ura3Δ0 leu2Δ0 his3Δ1 lys2Δ0 met15Δ0 can1Δ0::LEU2-MFA1pr::HIS3 hrr25-ts::URA3*, *pGBDC1-HRR25::TRP1* |
| SGY1168b | *MAT***a** | *ura3Δ0 leu2Δ0 his3Δ1 lys2Δ0 met15Δ0 can1Δ0::LEU2-MFA1pr::HIS3 hrr25-ts::URA3*, *pGBDC1-HRR25::TRP1* |
| SGY1428 | *MAT***a** | *trp1-901 leu2-3, 112 ura3-52 his3-200 gal4*Δ *gal80*Δ *Met2::GAL7-lacZ LYS2::GAL1-HIS3 GAL2-ADE2 pGBDC1-HRR25:: TRP1 + pGAD-C1::LEU2* |

**Table S2.** List of bacterial strains used in this study.

| **Strain** | **Genotype** |
| --- | --- |
| KC8 | *E.coli KC8 (US 15) Nx 1486 M+ K-12 leuB-600 trpC 9830 PyrF::Tn-5 hisB 463 del lacx74 Str A galU gal K* |
| DH5alpha | *fhuA2* Δ*(argF-lacZ)U169 phoA glnV44 Φ80* Δ*(lacZ)M15 gyrA96 recA1 relA1 endA1 thi-1 hsdR17* |

**Table S3.** List of plasmids used in this study.

| **Plasmid name** | **Source** |
| --- | --- |
| pGBDC1 | (James et al., 1996) |
| pGADC1 | (James et al., 1996) |
| pGBDC1+*HRR25* | This study |
| pAD-Chl4/pBD-Mcm19 | Ghosh et al., 2001 |

***Table S4. List of primers used in this study.***

| **Primer name** | **Description** | **Sequence** |
| --- | --- | --- |
| MA53 | Cloning of *HRR25* in pGBD vector (with *BamH*I site) | 5’ gagacgcggatcatggacttaagagtaggaag 3’ |
| MA54 | Cloning of *HRR25* in pGBD vector (*Pst*I site) | 5’ gagaaaaactgcagttacaaccaaattgactggc 3’ |
